# CellBin: a highly accurate single-cell gene expression processing pipeline for high-resolution spatial transcriptomics

**DOI:** 10.1101/2023.02.28.530414

**Authors:** Mei Li, Huanlin Liu, Qiang Kang, Shuangsang Fang, Min Li, Jiajun Zhang, Fei Teng, Dan Wang, Weixuan Cen, Zepeng Li, Ning Feng, Jing Guo, Qiqi He, Leying Wang, Tiantong Zheng, Shengkang Li, Yinqi Bai, Min Xie, Yong Bai, Sha Liao, Ao Chen, Yong Zhang, Susanne Brix, Xun Xu, Yuxiang Li

## Abstract

**Background:** Owing to recent advances in resolution and field-of-view, spatially resolved transcriptomics sequencing, such as Stereo-seq, has emerged as a cutting-edge technology for the interpretation of large tissues at the single-cell level. To generate accurate single-cell spatial gene expression profiles from high-resolution spatial omics data, a powerful computational tool is required.

**Findings:** We present CellBin, an image-facilitated one-stop pipeline for high-resolution and large field-of-view spatial transcriptomic data of Stereo-seq. CellBin provides a comprehensive and systematic platform for generating high-confidence single-cell spatial gene expression profiles, which specifically includes image stitching, image registration, tissue segmentation, nuclei segmentation and molecule labeling. CellBin is user-friendly and does not require a specific level of omics and image analysis expertise.

**Conclusions:** During image stitching and molecule labeling, CellBin delivers better-performing algorithms to reduce stitching error and time, in addition to improving the signal-to-noise ratio of single-cell gene expression data, in comparison with existing methods. Additionally, CellBin has been shown to obtain highly accurate single-cell spatial data using mouse brain tissue, which facilitated clustering and annotation.

## Introduction

Spatially resolved technology generates comprehensive data regarding the distribution of molecules that can be used to identify the location and function of cells within tissues, which helps to broaden our understanding of organ development [1], tumor heterogeneity [2] and cancer evolution [3,4]. This rapidly developing field is focused on obtaining detailed molecular information at single-cell resolution, as well as spatial information per molecule of tissue in a large field-of-view [5]. Single-cell resolution technologies [6–8], make it possible to explore spatial omics data at the single-cell, or even subcellular level. When combined with large field-of-view technologies [6], it allows for generation of 3D maps representing biological functions within single cells at the organ level.

Our previous work developed the high-resolution and large field-of-view spatially resolved technology, Stereo-seq, which enabled whole-organ sequencing of most tissues in model organisms (e.g., mouse brain (1cm×1cm) and mouse embryo (1cm×2cm)) [6]. The Stereo-seq technology generates two main outputs: spot-level gene expression data and high-content staining tissue images. The former is based upon detection of unique molecular identifiers captured by each DNA nanoball (DNB) on the sequencing chip while the latter provides detailed image-based (visual) information about the cellular matrix within tissue samples. Solutions that merge the two to obtain accurate spatial gene expression profiles at single-cell level in large field-of view settings will advance spatially resolved technologies and be an important steppingstone for downstream analyses.

Although spatially resolved technologies vary in resolution and field-of-view, one common feature is their ability to produce traditional high-content tissue images using dyes such as fluorescence, hematoxylin-eosin (H&E) or 4,6-diamidino-2-phenylindole (DAPI) for cellular or nucleus staining. The generated images provide important information regarding tissue and cell morphology, however, two main difficulties exist: insufficient precision in image mosaics and inaccuracy of molecule labeling.

The first difficulty is limited by the small imaging area, mechanical tolerance, and imperfect calibration of microscopes (e.g., 10×imaging lens, ∼1mm×1mm/tile), which makes them sufficient for visual inspection of tissue images, but not accurate enough for quantitative single-cell analytical approaches. For example, a 1cm×1cm object generates approximately 10×10 image tiles, while a 3cm×3cm object generates thousands of image tiles. Existing image stitching methods, such as ASHLAR [9] and MIST [10], can meet most stitching accuracy requirements. However, stitching misalignment is a major issue, and it is difficult to collect evaluation datasets to improve the accuracy of image mosaics in a large field-of-view, which is an obstacle in expanding their application.

The second difficulty, which is the key to achieving high-precision single-cell resolution, is classifying the transcripts and other molecules into the owner cell, a procedure termed “molecule labeling”. However, molecule labeling is not an easy task due to the low efficiency of molecule capture and the problem of diffusion of molecules outside cells during wet-lab procedures, which greatly affects the correct annotation of transcripts to the corresponding cell. Recent technology has considered this issue, such as 10X Xenium [11], which includes more transcripts by extending the distance outside the nucleus (15μm). Since the diameter of immune, stromal, and tumor cells varies, it is however challenging to obtain accurate single-cell data using such a generalized model that introduces varying levels of noise depending on the cell type. Some available methods, including Baysor [12] and Pixel-seq [13], require high-quality data as input, such as the number of captured molecules and their density distribution, which may not be applicable to most sequencing-based spatial omics data. Furthermore, existing spatial data analysis frameworks, such as Seurat [14], RCTD [15], Cell2location [16], Squidpy [17], and Spateo [18], only focus on the downstream analysis tasks or one specific aspect of the spatial data analysis or consider only cell nuclei segmentation.

Here we present CellBin, the first one-stop pipeline to generate single-cell spatial gene expression profiles for whole transcriptome based on high-resolution and large field-of-view spatially resolved sequencing. For image stitching, CellBin employs multiple Fast Fourier Transform (FFT) [19] weighted stitching algorithm, named MFWS, to reduce stitching errors in large-field-of-view datasets. The morphological tissue image stitching procedure of CellBin is accurate and reliable for single-cell identification and is flexible and convenient in terms of run time. Its high-precision stitching algorithm is useful for the correction of stitching to almost subcellular precision. For molecule labeling, CellBin employs the cell morphology and Gaussian Mixture Model (GMM) [20] algorithm, named MLCG, to increase the signal-to-noise ratio of the single-cell spatial gene expression profile, which yields more reliable analysis of cell clustering and annotation. CellBin has been successfully applied to datasets of various organs (such as brain, heart, embryo, artery, testis, kidney, liver and lymph) from different organisms (such as *Homo sapiens*, *Mus musculus*, *Macaca mulatta* and *Leporidae*). The molecule labeling of CellBin is also applicable to the datasets from multiple platforms other than Stereo-seq when the nuclei mask and spatial gene expression data are given. CellBin’s rich documentation in the form of a functional application programming interface, examples, and tutorial workflows, is easy to navigate and accessible to both experienced developers and beginner analysts. CellBin has the potential to serve as a bridge between the fields of image analysis and molecular omics, providing a foundation for the development of next generation computational methods for spatially resolved technologies.

## Results

### Overview of CellBin

CellBin is an image-facilitated one-stop pipeline for Stereo-seq transcriptomic data. CellBin combines large field-of-view tissue images and high-resolution spatial gene expression data to obtain high-confidence single-cell spatial gene expression profiles. CellBin provides a straightforward and systematic platform, which includes image stitching, image registration, tissue segmentation, nuclei segmentation and molecule labeling (Fig. 1a). We use MFWS for image stitching, which is based on frequency domain information and allows the accurate stitching of high-resolution images in a wide field-of-view, in addition to improvements in cell dislocation errors caused by inaccurate image stitching and low efficiency caused by the huge number of image tiles (Fig. 1b). We transform the spatial gene expression data into a map. The stitched image is registered with the spatial gene expression map based on “track lines” (marker lines designed on Stereo-seq chip), which includes translation, scaling, flipping and rotation (Fig. 1c). We train the deep learning models for tissue segmentation (Fig. 1d) and nuclei segmentation (Fig. 1e) from the registered image. We enable molecule labeling though MLCG, which applies GMM to fit the molecules within nuclei mask for accurate assignment of surrounding molecules to the most probable cell, obtaining a gene expression profile at the single-cell level (Fig. 1f).

**Figure 1:**
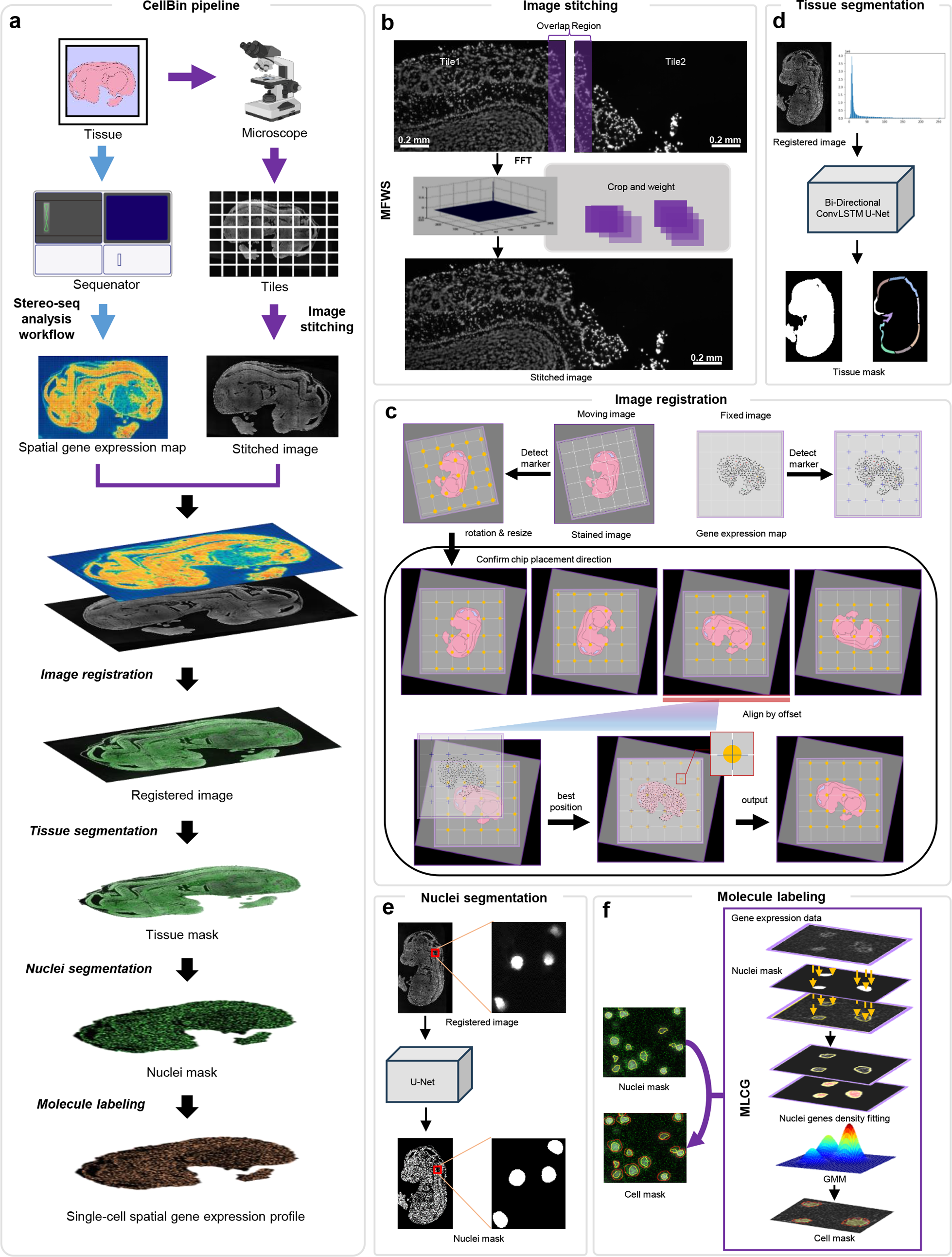
The CellBin workflow and applied algorithms. **a.** Schematic overview of the CellBin pipeline. The spatial gene expression data and the morphological (nuclei-stained) image of tissue are obtained using spatial transcriptomics technology and microscopy, respectively. The image tiles obtained by microscopy are stitched together to generate a large mosaic image of the whole tissue, the spatial gene expression data is transformed to a map, and the stitched image and spatial gene expression map are registered. Tissue and nuclei segmentations are performed on the registered image to obtain the tissue mask and nuclei mask. Molecule labeling is adopted to obtain single-cell spatial gene expression profile. **b.** Cutting the overlap regions of the adjacent image tiles, and calculating the spectral information based on the Fast Fourier Transform (FFT) algorithm. Weighted spectral information of the overlap cuts pairs to take the maximum spectral position as the offset of the adjacent image tiles. The image tiles are stitched using the offset value. **c.** Registration with spatial gene expression map and stitched image based on “track lines”. The spatial gene expression map is the fixed image, and the stitched image is the moving one. We detect the “track lines” on two images, the “track lines” on the spatial gene expression map represent the template, for the stitched image, we calculate the scale and rotation parameters using the “track lines”. We then use the morphological features to get a rough registration of the offset, flip, and 90° rotation. “Track lines” are used to fine-tune registration. **d.** Bi-Directional ConvLSTM U-Net is used to obtain the tissue mask on registered image. **e.** U-Net is used to obtain the nuclei mask on registered image. **f.** Molecules within the nuclear boundary are assigned to a given cell, while molecules outside the nucleus are labeled to the cell that gains the largest probability from Gaussian Mixture Model fitting.

We provide a fixed set of hyperparameters for CellBin that are suitable for most Stereo-seq datasets. This helps prevent users from feeling overwhelmed by the numerous hyperparameters involved in the all-in-one process. We examine the application of CellBin on various tissue datasets and compare it with other state-of-the-art methods. We validate the efficiency and effectiveness of MFWS on Stereo-seq and public datasets, comparing it with ASHLAR [9] and MIST [10]. The accuracy of the generated single-cell spatial gene expression profile is evaluated on a mouse olfactory bulb dataset, confirming the efficacy of MLCG. We analyze the cortex structure in the generated single-cell spatial gene expression profile on a mouse brain dataset, comparing the results with those obtained using the Bin*X* method (Bin*X* means a bin with *X* × *X* of DNBs [6]). As there is currently no image-based one-stop pipeline like CellBin available for Stereo-seq data, we compare CellBin with SCS [21], which generates spatial gene expression profiles based on nuclei segmentation masks but lacks image stitching and registration capabilities, and Baysor [12], which can generate spatial gene expression profiles without using staining images. The main experimental results are shown below, and additional experimental results can be found in Supplementary materials and Supplementary Fig. S1.

### CellBin processes high-precision morphological tissue images from image tiles

MFWS of CellBin takes a folder of all image tiles obtained by microscopy and information files as the input, and outputs a stitched mosaic image of whole tissue. The datasets from different field-of-views (4 Stereo-seq mouse brain datasets with the chip sizes of 1cm×1cm, 1cm×2cm, 2cm×2cm and 2cm×3cm respectively and a public dataset [22], and their grid sizes (11,9), (15,21), (25,21), (23,29) and (10,10), respectively) are collected. For each dataset, the standards are designed (Supplementary materials) to calculate the relative offset error between each two adjacent image tiles and the absolute offset error of the entire stitched image in the stitching results of different methods. The relative offset errors are statistically analyzed by Wilcoxon signed rank test [23] to examine the significant differences. The runtime to obtain the absolute offset error for each method is recorded.

Processing a morphological image from a tissue slide requires the stitching of an array of multiple image tiles generated by microscopy. Microscopes can automatically capture the image tiles of a tissue one by one and stitch the image tiles together using the built-in stitching method. However, overlapping areas between adjacent image tiles generated during microscope movement may be imprecise due to mechanical tolerance and imperfect calibration. Such stitching errors are common in mosaic images and need to be removed if the goal is to achieve single-cell resolution in spatial omics applications. As an example, a mouse brain mosaic image on a chip with a size of ∼2cm×2cm and a tile size of 2424pixels×2031pixels are displayed (Fig. 2a), corresponding to a physical size of ∼1.2mm×1.0mm. To visualize seams and stitching errors, we select two adjacent image tiles from the image (Fig. 2b). For accurate stitching results, the lower part of tile 1 (dotted area) and the upper part of tile 2 (dotted area) should be accurately overlaid. Shadows that represent inaccurate overlap of cells can be easily seen in the stitching results produced by the microscope (Fig. 2c, left), but MFWS of CellBin is able to stitch the image tiles accurately, resulting in the absence of shadows (Fig. 2c, right). The stitching results of two image tiles within non-tissue areas (full-colored part) where individual “track lines” can be clearly seen using auto-contrast adjustment as also shown (Fig. 2c). The applied Stereo-seq chip contains periodic “track lines” with a size of 1500 nm (∼3 pixels), which is apparent from the image (Fig. 2c, f). We use a collection of data to evaluate the accuracy and efficiency of existing stitching algorithms as compared with MFWS.

**Figure 2:**
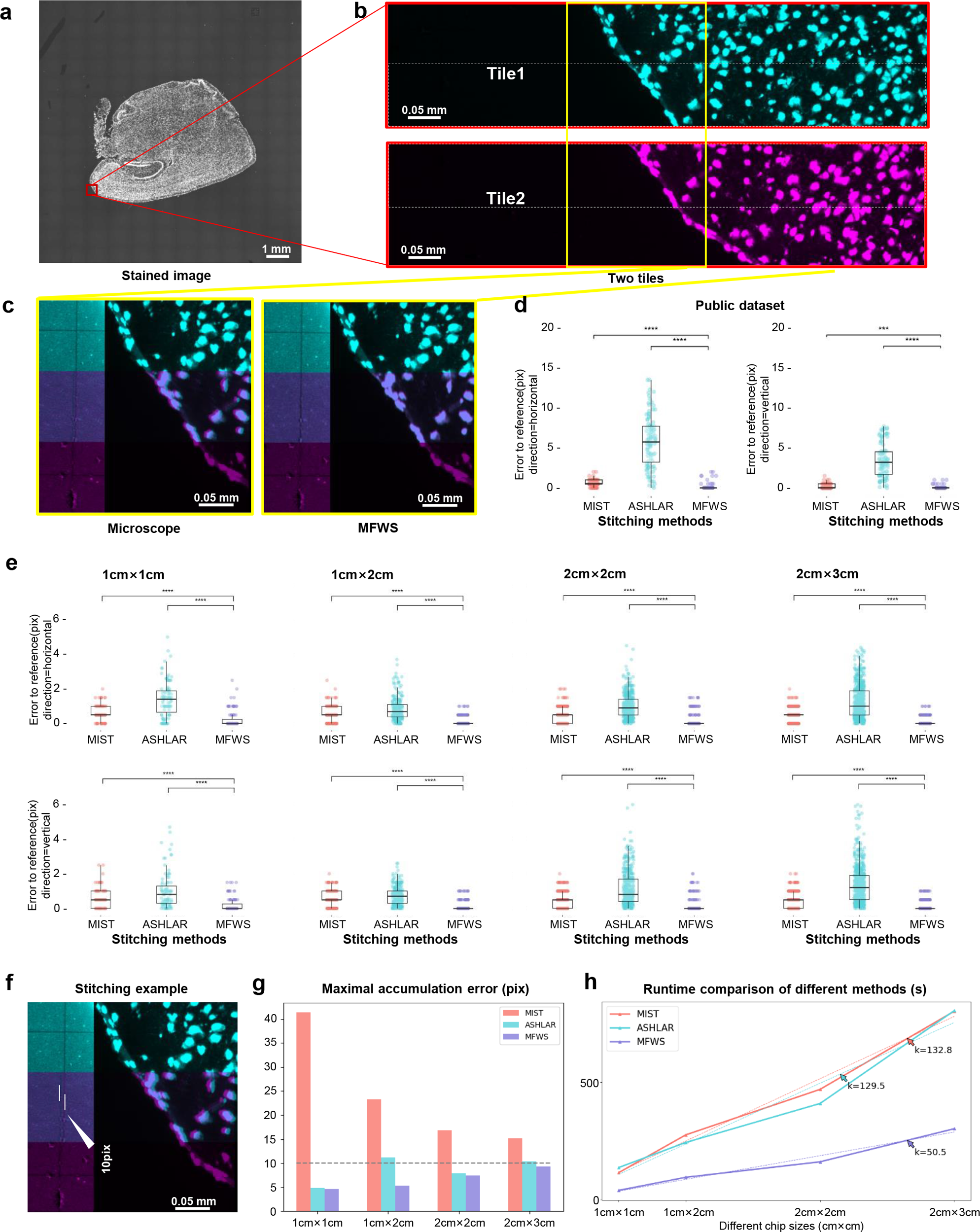
CellBin includes a high-precision mosaic method for image stitching, enabling large field-of-view morphological image analysis at single-cell spatial accuracy. **a.** Mosaic image of mouse brain tissue on a 2cm×2cm chip. **b.** The red box indicates the edge of the tissue (containing cells and non-tissue), and two neighboring image tiles (in the vertical direction) are shown in different colors. The part that needs to be overlaid by stitching is the lower part of tile 1 (within the dotted area) and the upper part of tile 2 (within the dotted area). **c.** The stitching results for the two image tiles (yellow box in **b**) using the stitching method built into the microscope (left) and MFWS (right). In each sub-figure, the left part shows the overlap of cells after stitching and the right part shows the “track lines” in the non-tissue area. The contrast of the background was increased to clearly show the “track lines” incorporated into the Stereo-seq chip. **d.** Comparison of the relative errors produced by MIST, ASHLAR, and MFWS on a public dataset. **e.** Comparison of the relative errors produced by MIST, ASHLAR, and MFWS on Stereo-seq mouse brain datasets, analyzed using different chip sizes from 1cm×1cm to 2cm×3cm, corresponding to number of image tiles from 11×9 to 23×29. The evaluation is based on the ground truth calculated using the “track lines” on the Stereo-seq chip. **f.** An example of the stitching error in pixels in the zoomed-in image. A dislocation of 10-pixels roughly corresponds to half a cell. **g.** Bar graph illustrating the maximal accumulation of stitching errors produced by MIST, ASHLAR, and MFWS on the datasets from **e**. **h.** Line graph displaying the run time of MIST, ASHLAR, and MFWS on Stereo-seq mouse brain datasets.

We apply MFWS to the public dataset, and the results show that the relative offset errors of MFWS are comparable with those of MIST [10], both being concentrated within 5 pixels, while ASHLAR [9] has larger offsets (>10-pixels) (Fig. 2d). Moreover, the offset error distribution is much more concentrated by MFWS. Next, we calculate the relative and absolute offset errors on 4 Stereo-seq mouse brain datasets. MFWS is shown to perform significantly better than ASHLAR [9] and MIST [10] with respect to the relative offset errors for all image size combinations (Fig. 2e). A 10-pixel (∼5μm) dislocation roughly corresponds to half a cell (Fig. 2f). In all the datasets, the errors generated by MFWS remain within a maximum of 10-pixels in the tissue area, while the other methods produce a shift greater than 10-pixels, which may misplace the cell (Fig. 2g). As the number of image tiles increases, the run time becomes a significant factor for stitching algorithms. The run time of MFWS is significantly shorter than that of the other methods, mostly due to the embedded spectral calculation based on FFT (Fig. 2h).

### CellBin generates highly accurate single-cell gene expression profiles of spatial omics datasets

Using CellBin, we first obtain a nuclei mask and a cell mask, and then output their single-cell spatial gene expression profiles. To demonstrate the improvement in transcript assignment to cells by using the cell mask generated by CellBin, we here compare the gene expression profile output using each of these masks on Stereo-seq data generated of mouse olfactory bulb, involving analysis of spatial gene expression data containing 37,288,344 molecules and 143 image tiles. The generated profiles are input into Stereopy (v6.0) [24] (a downstream analysis tool) for analysis. The silhouette coefficient [25] is used to evaluate clustering results. The moran’s I (calculating by Scanpy [26] package) is used to evaluate spatial correlation. The details of the data analysis process and evaluation metrics are provided in Supplementary materials.

After obtaining the nuclei mask, the molecules located inside the nucleus are assigned to the corresponding cell, and MLCG of CellBin fits the molecular density distribution to each cell nucleus and re-labels the molecules outside the nucleus to finally generate a cell mask (Fig. 3a). By applying the cell mask, the uniquely expressed genes and total gene counts in a single cell increased by approximately 2.34 and 2.56 times, respectively, as compared to the nuclei mask alone (Fig. 3b). Utilization of the cell mask provides better overall clustering (Fig. 3c), and its silhouette coefficient is higher than that of the nuclei mask. We manually annotate the cell types by comparing the differentially expressed genes in each cluster with the marker genes of cell types inferred by a reference dataset [27]. Of note, there are fewer scattered points and a more concentrated distribution in the spatial position of the astrocyte layer (Moran’s I: 0.40 vs. 0.30) and dopaminergic neuron layer (Moran’s I: 0.52 vs. 0.37) after molecule labeling using the cell mask (Fig. 3c). We further explore the marker gene distribution and expression in the annotated cell types, and all the marker genes provided by the reference dataset [27] display higher expression using the cell mask as compared with the nuclei mask (Fig. 3d). We compare subtypes of granule cells discovered using the nuclei mask vs. the cell mask. The nuclei mask is able to identify two subtypes of granule cells: granule cell 0 and granule cell 3, while the cell mask enables identification of three granule cell subtypes: granule cell 0, granule cell 1, and granule cell 3. We explore the expression of the granule cell 1 subtype-marker genes Syt1, Scg2, and Cplx1, among all granule cells. There is no obvious difference in expression when using nuclei mask, while higher expression is found in granule cell 1 as compared with the other two subtypes when using cell mask (Fig. 3e). This supports that the cell mask captures a higher number of transcript signals from rare cells, facilitating better annotation and thereby enhances single-cell spatial resolution.

**Figure 3:**
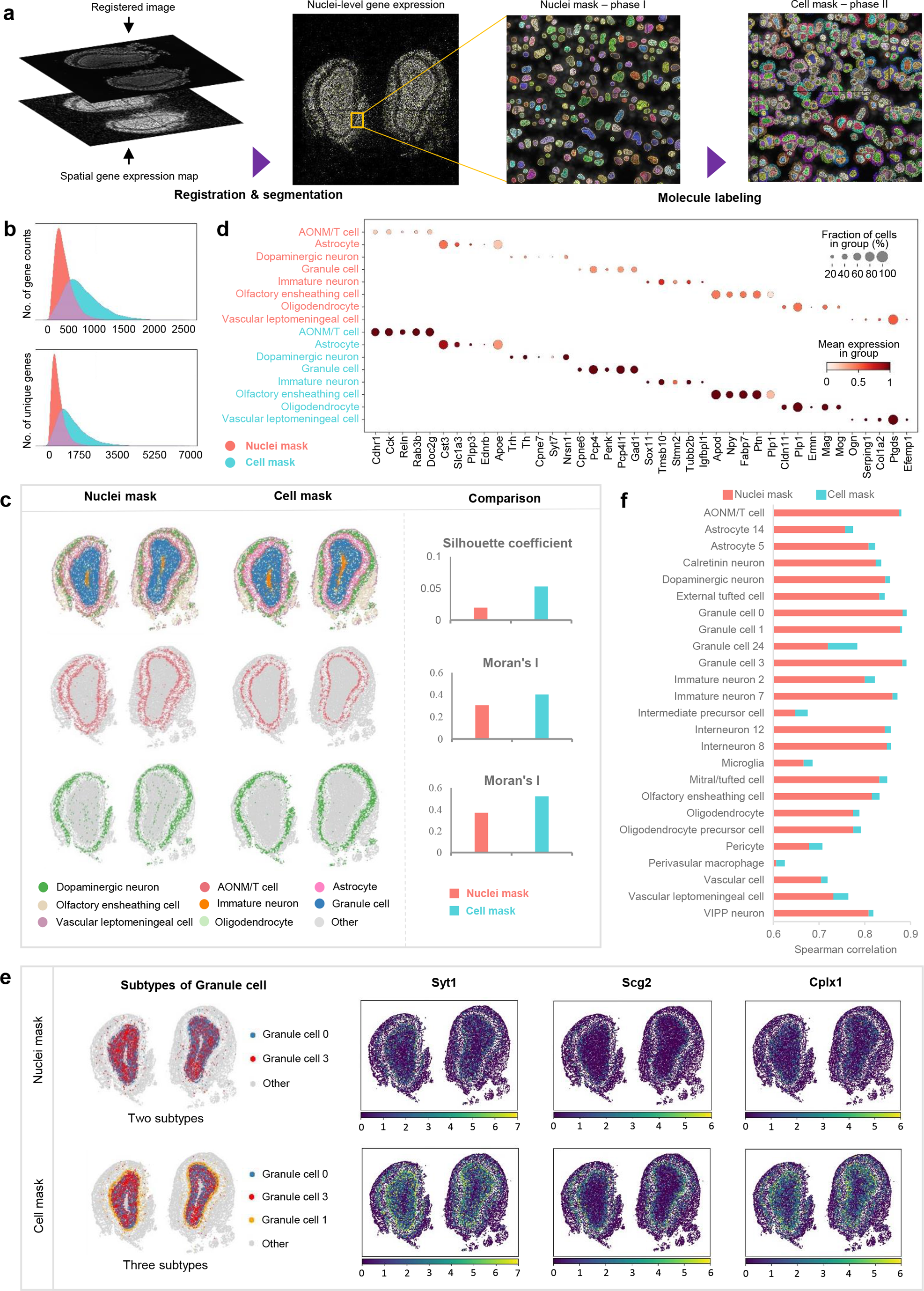
CellBin provides single-cell spatial data with a higher signal-to-noise ratio that facilitates finer cell clustering and annotation of the mouse olfactory bulb. **a**. Flowchart of MLCG processing of Stereo-seq mouse olfactory bulb dataset. **b**. Comparison of the number of uniquely expressed genes and the total gene count per cell using nuclei mask vs. cell mask on Stereo-seq mouse olfactory bulb dataset. Top: density plot of the total gene counts per cell. Bottom: density plot of the number of uniquely expressed genes per cell. **c**. Comparison of the clustering results generated using the “Leiden” algorithm on nuclei mask vs. cell mask. Top row, left: spatial clustering results of mouse olfactory bulb data generated using nuclei mask vs. cell mask; right: comparison of unique cells estimated by Silhouette coefficient. Middle and bottom rows, left: spatial distribution of cells in the inferred clusters of astrocytes and dopaminergic neurons generated using nuclei mask vs. cell mask; right: comparison of the spatial autocorrelation estimated by Moran’s I. **d**. Comparison of marker gene expression in the main cellular clusters inferred by the “Leiden” algorithm generated using nuclei mask vs. cell mask. **e.** Comparison of cellular subtype identification using nuclei mask vs. cell mask. First column: subtypes of granule cells identified using nuclei mask vs. cell mask. Other columns: gene expression heat maps of Syt1, Scg2, and Cplx1 respectively, which have been reported to be marker genes for granule cell 1, using nuclei mask vs. cell mask. **f**. Spearman correlation between gene expression in similar cells in tissue slides of the mouse olfactory bulb using nuclei mask vs. cell mask to define the gene expression in single cells and a single-cell reference dataset of the mouse olfactory bulb. **c**, **d**, **f**: AONM/T cell: anterior olfactory nucleus mitral/tufted cell, VIPP neuron: vasoactive intestinal peptide positive neuron.

We estimate and compare the similarity of spatial gene expression between a reference single-cell dataset of the mouse olfactory bulb [27] and our Stereo-seq based mouse olfactory bulb dataset using the CellBin nuclei mask vs. the cell mask to generate single-cell expression profiles. Tissue cells are auto-annotated by Tangram [28] and correlation analysis is performed between the expression matrixes of single cells using each of the masks and the single-cell sequencing expression profile of the reference [27]. The Spearman correlation coefficient using annotations based on the cell mask is higher (0.4% to 8.9% better) than that based on the nuclei mask. Overall, the single-cell expression matrix generated by the cell mask shows a higher correlation with the single-cell reference (Fig. 3f), indicating that the cell mask provides a spatial single cell-level gene expression profile closer to that generated by single-cell sequencing.

### CellBin enables dissection of the structural composition of mouse brain cortex data at single-cell resolution

Non-cell-based binning methods where adjacent tissue regions are divided into regions (bins) of specified sizes, are sometimes used in spatial transcriptomics analysis pipelines. To compare the CellBin cell mask output to the Bin*X* method, we here apply a Stereo-seq mouse brain dataset containing 131,990,020 molecules and 117 image tiles, where image tiles where split using Bin100, Bin50, Bin20. The generated profiles are input into Stereopy (v6.0) [24] for analysis, and moredetails are provided in Supplementary materials. We export the spatial gene expression map, nuclei-stained image, cell mask and Bin20 outlines to visually demonstrate the segmentation effect of the two methods. We also compare the abilities of CellBin and the Bin approach to reconstruct known cellular regions in the mouse brain.

The clustering of data generated by CellBin vs. Bin100, Bin50, and Bin20 is shown (Fig. 4a). It appears that splitting using Bin100 results in difficulties in identifying several important areas in the tissue, such as the brain hub and hippocampus. For Bin50, the tissue cortex and blood cells are poorly identified. Both Bin20 and CellBin can identify important areas in the tissue, but CellBin obtains the highest silhouette coefficient in evaluating the clustering results using several methods (Fig. 4b). We calculate the uniquely expressed genes and total gene counts of CellBin vs. the differently sized bins (Fig. 4c). The visualization shows that Bin20 more often divides a single cell into two or more cells, while CellBin more accurately divides the cell area and is consistent with the actual cell distribution in the tissue (Fig. 4d). Bin20 results in splitting of approximately 90% of the cells, with only ∼10% of the nuclei being completely covered in the nuclei-stained image, while only ∼2% of the cells are split into two or more cells using CellBin (Fig. 4e). The resulting single-cell data derived from cell identification using Bin20 and CellBin were individually annotated using Spatial-ID [29] (Fig. 4f) with adolescent mouse brain as a reference [30]. Bin20 annotates 29 different cell types, while CellBin can annotate 37 different cell types. Within the ACTE series, both Bin20 and CellBin annotate 2 subtypes. In the MEINH series, Bin20 annotates 2 subtypes, while CellBin annotates 3 subtypes. Within the TEGLU series, Bin20 and CellBin annotate 10 and 13 subtypes, respectively. In TEINH series, Bin20 annotates 5 subtypes and CellBin annotates 4 subtypes, which is the only case where fewer subtypes were annotated by CellBin. CellBin also annotates 2 subtypes within the ACNT series and 4 subtypes within the TEINH series. When zooming in on the cortical area (Fig. 4g), it appears that the staining position of CellBin and the nucleus are better aligned than seen for Bin20, and the cell state and tissue structure are more in line with the actual brain tissue map. Moreover, in the gear gyrus and cortical regions of the mouse brain, CellBin performs better than Bin20 in matching the annotation results to the Allen mouse brain dataset (Fig. 4h). Although Bin20 is capable of annotating the gear gyrus and cortex to align with the location in the Allen mouse brain atlas, the division between layers is not as accurate. Expression of marker genes in the gear gyrus (DGGRC2) and cortex (TEGLU3, TEGLU4, TEGLU7, and TEGLU8) is fully captured using the CellBin algorithm, which is not the case when using Bin20 (Fig. 4i). Overall, CellBin provides single-cell spatial gene expression profiles with a higher signal-to-noise ratio compared to approaches based on differently sized bins. Moreover, the profile generated using CellBin aligns well with cellular annotation within different brain regions and can provide a valuable reference for studies of cellular interaction networks in health and disease.

**Figure 4:**
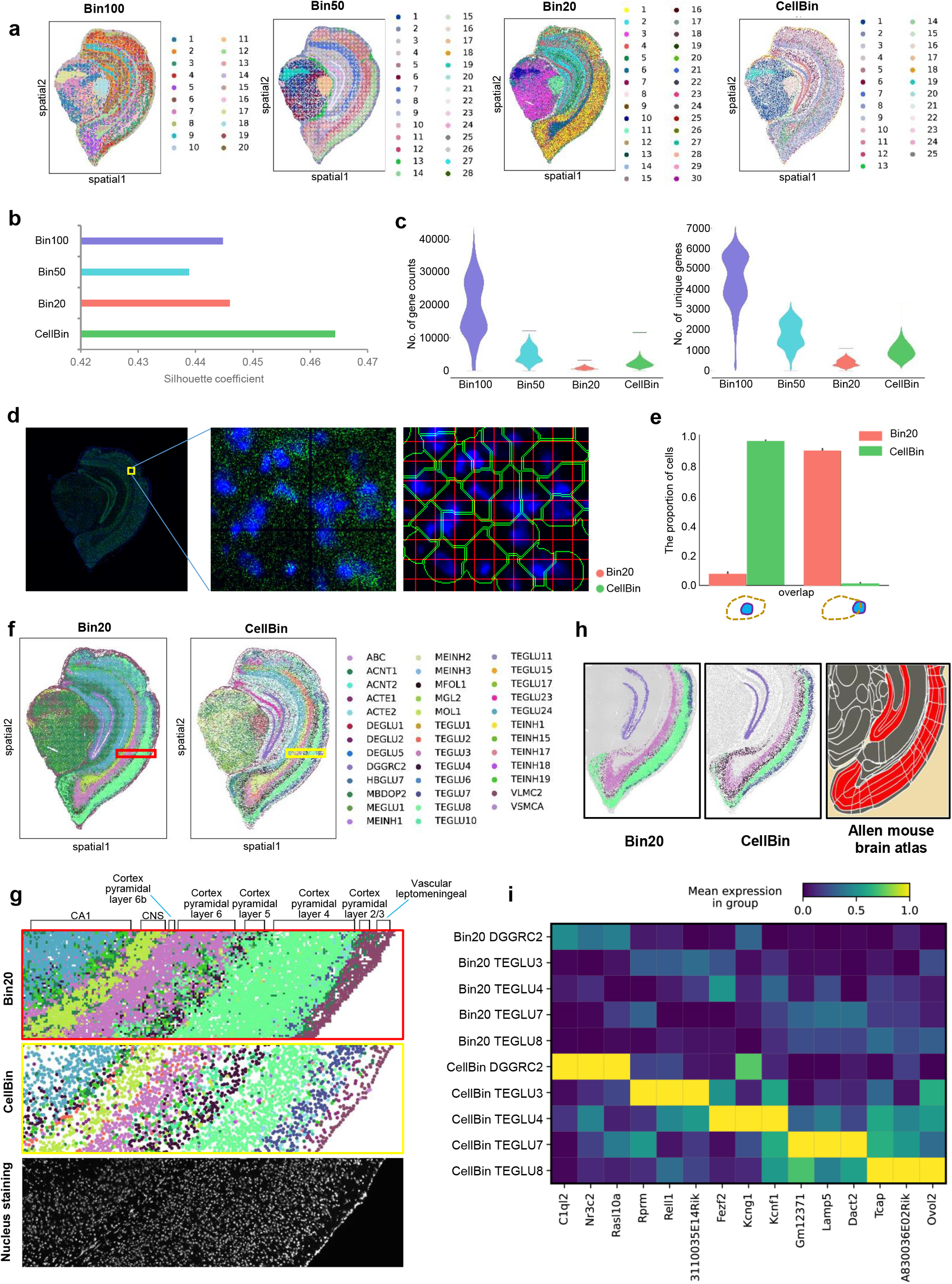
Clustering and annotation of mouse brain tissue by CellBin vs. differently sized bins. **a.** Identification of gene expression clusters using differently sized bins vs. CellBin on Stereo-seq mouse brain dataset. **b.** Silhouette coefficient evaluation of clustering performance. **c.** Violin plots displaying the total gene counts and number of uniquely expressed genes per cell captured using differently sized bins vs. CellBin. **d.** Comparison of the cell boundary generated using Bin20 and CellBin. The first image shows the spatial gene expression map and nuclei-stained image, which are merged in green and blue, respectively. The second image displays an enlarged map of the local area from the first image. The third image is a comparison of the cell boundaries obtained using Bin20 (red) vs. CellBin (green). **e.** Comparison of the proportion of cell nuclei covered or intersected using Bin20 vs. CellBin. **f.** Comparison of end-to-end annotation results of Bin20 vs. CellBin. Colors represent the same cell type in the Bin20 and CellBin annotation charts. **g.** The enlarged local comparison map of the cortex shows the spatial distribution and boundary information of different cortical cells. **h.** Comparison of cell annotation based on Bin20 vs. CellBin using the Allen mouse brain atlas as a reference. The spatial distribution of the gear gyrus (DGGRC2, black) and the cortex (TEGLU3, TEGLU4, TEGLU7, and TEGLU8) is shown. **i.** Heat map displaying the consistency of marker gene expression within the different substructures in **h** (gear gyrus (DGGRC2) and cortex (TEGLU3, TEGLU4, TEGLU7, and TEGLU8) layers)). The upper part: results of Bin20, lower part: results of CellBin.

## Discussions

In the present study, we demonstrate that auxiliary staining can be extremely valuable to resolve difficult cases. The molecule labeling or cell segmentation results will likely be improved using better staining, such as labeling of outer membranes, and better techniques for segmenting such images. Additionally, better photo capturing using enhanced AI recognition models will advance our insight into biological traits and speed up the identification of features that enable us to distinguish between healthy and diseased tissue. It is noteworthy that the entire CellBin framework can be applied to generate single-cell gene expression for Stereo-seq, and most of its modules are also suitable for other spatial transcriptomics technology.

CellBin has quality requirements for the input data. For Stereo-seq transcriptomic data, the gene expression counts in each Bin200 should not be less than 5000. For the image tiles taken by the microscope, the “track lines” clarity is required to pass our image quality control. The registration of CellBin involves aligning the data of two different modalities, i.e. spatial gene expression data and staining image. To ensure higher accuracy and stronger robustness in the registration process, the assistance of “track lines” is still indispensable and irreplaceable at present.

CellBin is equipped with an extensive testing suite, implemented in a robust continuous integration pipeline that contains a graphic user interface for manual image stitching, image registration, and molecule labeling. CellBin offers a rich documentation, including a functional application programming interface, examples, and tutorial workflows, making it easy to navigate and accessible for diverse developers and analysts. CellBin can be used on desktops, laptops, workstations, computer clusters or cloud computing platforms. To ensure the normal operation of CellBin, we recommend a hardware configuration of at least 8-core CPU and 16 GB of RAM. The use of GPU can greatly accelerate the tissue and nuclei segmentation steps. The amount of disk storage required depends on the input dataset. The information and timing records on more different tissue datasets are shown in Supplementary Table S1, which can assist the users in estimating the required time for their specific situations.

## Conclusions

There is still a long way to go before the potential of spatially resolved technology can be fully utilized, and continued improvement of both protocols and analytical methods for processing spatial omics data is required. Here, we present CellBin, an image-facilitated cell segmentation framework for high-resolution and large-field-of-view spatial omics that provides a complete and systematic platform for obtaining high-confidence single-cell spatial data. The implemented morphological tissue image stitching procedure proved to be accurate and reliable for single-cell identification and was flexible and convenient in terms of run time. The high-precision stitching method based on FFT and was found to be useful for the correction of stitching to almost subcellular precision. We implemented molecule labeling based on GMM to generate spatial single-cell gene expression profile with a higher signal-to-noise ratio, which yielded more reliable analysis of cell clustering and end-to-end annotation. Moreover, we optimized the processing modules used for image registration to include “track lines” on the chip, nuclei segmentation, and tissue segmentation.

The goal here is to develop a general pipeline for profile generation and single-cell level analysis of spatial omics data. The data presented herein are based on spatial transcriptomics alone. In future studies, we will combine spatial information with additional biological characteristics to integrate multi-omics information at the single-cell spatial level, improve the signal-to-noise ratio, and develop interfaces and extensions that support user-defined data and methods.

## Methods

### Image stitching

The image tiles are stitched into a large mosaic image of the whole tissue by MFWS (Fig. 1b). The file name of each image tile needs to reflect the row and column, such as “0000_0001.tif” that reflects a tile at row 0 and column 1. Firstly, we calculate the actual overlap value with the neighbors in the horizontal and vertical directions for each image tile. A pair of image tiles in the vertical direction is taken as an example. (i) The overlapping regions of adjacent image tiles are marked as *A_f_* and *A_m_* respectively, and they are divided into *N* sub-region images as marked {*f_i_*} and {*m_i_*} respectively. (ii) The FFT [19] algorithm is applied to obtain 2×*N* frequency spectra based on {*f_i_*} and {*m_i_*}. (iii) The sub-region images in *A_f_* and *A_m_* are paired, such as *f_i_* and *m_i_*, and *N* cross-power spectra are obtained using the formula for image cross-power calculation. Theoretically, any one of these spectra can reflect the true offset information. (iv) Due to the low signal-to-noise ratio of some sub-regions in the previous step, the calculated value differs greatly from the actual value. The overall accuracy of the algorithm can be significantly improved by weighted enhancement of partial blocks and reduction of residual sub-blocks. Our weighting coefficient is based on the peak value of the mutual power spectrum because experiments show that the greater the peak value, the higher the precision and reliability of the overlap value. A unique cross-power spectrum is obtained after the weighted superposition of *N* spectra. Then, the corresponding spatial domain correlation graph can be obtained through Inverse FFT. The coordinate of the peak value represents the required offset value. (v) Carrying out the above steps for each pair of adjacent image tiles to generate the offset matrices *O_H_* and *O_V_*. Secondly, transformation of local coordinates to stitching coordinates is required, since image stitching demands the unique coordinates of each tile in the reference system, i.e., the global coordinate matrix *L*. In the *O_H_* and *O_V_* matrixes, offsets corresponding to low-confidence values are not credible that need to be eliminated. However, it leads to the existence of single or multiple connected domains. Therefore, when obtaining the coordinates, we first find the center of each connected domain, complete the splicing in the connected domain according to the relative relationship among the neighbors, and then use the experience value to fill multiple connected domains to complete the splicing of the entire image. Thirdly, according to the image tiles and the coordinate matrix *L*, the size of the mosaic image is obtained, the value is traversed, and the seam fusion is completed synchronously during the stitching process.

### Image registration

The spatial gene expression data is read and transformed into a map, in which the intensity value of each pixel is proportional to the total count of molecules of all genes expressed at position DNB. The stitched image is registered to the transformed map (Fig. 1c). Sequencing chips with sample tissue attached, imaging, and sequencing are designed with periodic “track lines” (horizontal and vertical) to assist base calling and image registration. The “track lines” are displayed as dark straight lines in both the stitched image and transformed map. Image registration is achieved using the transformed map as a template and performing an affine transformation on the stitched image. The “track lines” are detected in the transformed map and stitched image by line searching algorithms, and the intersections of the horizontal and vertical “track lines” are located. The scaling parameter is calculated by comparing the length of the line segments between intersections within the transformed map and stitched image. The rotation angle is defined as the difference between the horizontal “track lines” and the horizontal direction of the image coordinate system. For the offset between them, since the “track line” patterns are periodic, we first calculate the center of gravity of the tissue regions within the transformed map and stitched image, roughly match them together, and then match the “track lines” intersections to fine-tune the image registration.

### Tissue segmentation

The tissue can be segmented from the registered image. The tissue segmentation process consists of two steps: preprocessing and model inference (Fig. 1d). In addition, a self-scoring mechanism is added to optimize the processing time and eliminate the effort of manually checking the results. The histogram enhancement method is used to improve the separation of tissue and background during the preprocessing, and then, Bi-Directional ConvLSTM U-Net [31] is used to predict tissue masks during the model inference. Compared with the original U-Net [32], the following improvements are incorporated. (i) The convolution sequence is replaced with a densely connected convolution in the final step of the encoding path to encourage feature reuse. (ii) The feature maps extracted from the corresponding encoding path are combined with the previous decoding up-convolutional layer in a non-linear manner. (iii) The batch normalization is added to accelerate the convergence of the network. When dealing with a large amount of unstable data, the model cannot achieve ideal results with very complex or poor-quality tissue images, and the self-evaluation mechanism help to filter these potentially unsatisfactory results. When the registered image possesses a high gray value, we use the threshold value obtained in the histogram enhancement step as a reference value and assume that the pixel points higher than this value are valid pixels. The effective pixel densities of the tissue inside the mask and in the area surrounding the mask are calculated in sections, and the ratio of the two values is used as the basis for evaluation.

### Nuclei segmentation

The nuclei can be determined from the registered image due to nuclei staining. Nuclei segmentation consists of three steps (Fig. 1e). The first step is image preprocessing. The median filtering is employed to smooth the noise that may be present in the input image. To both enhance the cell and homogenize the image background, the output of the median filtering operation is input into the pixel-wise subtraction process. This step facilitates the relative conspicuity of nuclei against the background. The second step is segmentation model inference. We use U-Net [32] for segmenting the cells. Some optimizations are made to U-Net that, Residual U-Net [33] is used as a feature extractor and the Pyramid Squeeze Attention [34] module is used instead of the 3×3 convolution in the bottleneck as the basic feature-extracting unit of Residual U-Net, which ensures the model to pay more attention to the highlighted feature representation. The third step is mask post-processing. To obtain a more reliable segmentation, the mask generated by the segmentation model is input to the area filtering operation. Further, the opening operation (erode and dilatate) is applied to revise the cells shape and boundaries.

### Molecule labeling

The single-cell spatial gene expression profile can be obtained by combining the nuclei mask and spatial gene expression data (Fig. 1f). Firstly, the morphological tissue image is used to identify nuclear boundaries as described in nuclei segmentation. The specific transcripts that are contained within each nucleus are then assigned. A GMM [20] algorithm is used to estimate the probability of each transcript belonging to a given cell based on the nuclei mask. It is performed by modeling each cell as a GMM distribution, combining its spatial position, transcript count, and density. Simultaneously, we also estimate the probability scores of transcripts in the acellular region. In our model, the probability of assignment of a transcript to a given cell declines progressively with distance from the cell center but increases with transcript count and density. A mixture model is a probabilistic model that can be used to represent a population distribution with *K* sub-distributions. GMM can be regarded as a model composed of *K* single Gaussian models. Taking a single cell as an example, based on the result of nuclei segmentation, we can locate the spatial position of the nucleus. Firstly, GMM is used to fit the molecular distribution of the current cells. We expanded the nuclei mask boundary and captured as much as possible of the true distribution of data in cells rather than nuclei to develop the cell mask. According to the empirical value of the cell area, we finally determine the fitting range of 100pixels×100pixels. Subsequently, GMM is used to calculate the probability score of extracellular molecules within the fitting range. Finally, according to the maximum and minimum probability scores within the fitting range, the quartile is used as the adaptive threshold of the current cell. When the probability score is greater than the threshold, the molecule is re-divided into cells. In this way, molecules are assigned to their corresponding cell with high confidence.

## Availability of Source Code and Requirements

Project name: CellBin

Project homepage: https://github.com/STOmics/CellBin/tree/dev

Operating system(s): Windows or Linux

Programming language: Python

Other requirements: See https://github.com/STOmics/CellBin/blob/dev/requirements.txt for details

License: MIT License

RRID: SCR_025413

BiotoolsID: CellBin

## Data availability

The data that support the findings of this study have been deposited into Spatial Transcript Omics DataBase (STOmics DB) of China National GeneBank DataBase (CNGBdb) with accession number STT0000027: https://db.cngb.org/stomics/project/STT0000027.

## Addition Files

**Supplementary materials.** Stitching standard design, data analysis process, evaluation metrics, additional experiments and information of different tissue datasets.

**Supplementary Fig. S1.** CellBin can be applied to diverse tissue datasets and outperform other state-of-the-art methods. **a.** The clustering results of single-cell spatial gene expression profiles obtained by CellBin on diverse tissue datasets. **b.** The result comparison of distance and area errors obtained by MFWS, MIST and ASHLAR on a public dataset. **c.** The nuclei staining image of Subset 1 and result comparison of number of cells, number of clusters, silhouette coefficient, Moran’s I and running time obtained by CellBin, SCS and Baysor on Subset 1. **d.** The nuclei staining image of Subset 2 and result comparison of number of cells, number of clusters, silhouette coefficient, Moran’s I and running time obtained by CellBin, SCS and Baysor on Subset 2.

**Supplementary Table S1.** Timing values of CellBin on different tissue datasets.

## Authors’ contributions

Conceptualization: Mei Li, Huanlin Liu, and Shuangsang Fang.

Project administration and supervision: Susanne Brix, Xun Xu, Yong Zhang, and Yuxiang Li.

Soft development and implementation: Huanlin Liu, and Min Li.

Data collection, processing, and application: Mei Li, Huanlin Liu, Qiang Kang, Jiajun Zhang, Fei Teng, Weixuan Cen, Zepeng Li, Ning Feng, Jing Guo, Qiqi He, and Leying Wang.

Method comparisons: Huanlin Liu, and Qiang Kang.

Biological interpretation: Qiang Kang, Shuangsang Fang, Jiajun Zhang, and Fei Teng.

Data visualization: Tiantong Zheng and Shengkang Li.

Project coordination: Mei Li, Shuangsang Fang, Sha Liao, and Ao Chen.

Manuscript writing and figure generation: Mei Li, Huanlin Liu, Qiang Kang, Shuangsang Fang, Jiajun Zhang, Fei Teng, and Dan Wang.

Manuscript review: Susanne Brix, Yinqi Bai, Min Xie, and Yong Bai.

## Abbreviations

DNB: DNA nanoball
H&E: hematoxylin-eosin
DAPI: 4,6-diamidino-2-phenylindole
FFT: Fast Fourier Transform
MFWS: multiple Fast Fourier Transform weighted stitching algorithm
GMM: Gaussian Mixture Model
MLCG: cell morphology and Gaussian Mixture Model algorithm.

## Acknowledgements

This work is part of the “SpatioTemporal Omics Consortium” (STOC) paper package. A list of STOC members is available at http://sto-consortium.org. We would like to thank the MOTIC China Group, in addition to Bichao Chen, Canqiang Xu, Xiangxiang Liao, Kui Su, Zhonghan Deng, Yiwen Wu, Bohan Zhang, Chao Liu, Mengnan Cheng, Qing Zhou, Marija Ivanovic, Vladimir Kovacevic, Lei Han, Xing Guo, and Wei Peng for their help. This work was supported by Wuhan Supercomputing Center for the data training. We thank China National GeneBank for providing technical support. Dan Wang is supported by Zhuhai Basic and Applied Basic Research Foundation (2220004002717).

