## Supplementary materials for "CellBin: a highly accurate single-cell gene expression processing pipeline for high-resolution spatial transcriptomics"

### Stitching standard design, data analysis process, evaluation metrics, additional experiments and information of different tissue datasets

**The standard design in comparison experiment of image stitching methods.**

For each dataset, a standard is designed according to the size of each image tile and the translation parameter set during the microscope shooting, and the standard is fine-tuned manually to correct mechanical errors according to the overlap area between each two adjacent image tiles. These standards are used to calculate the relative offset error between adjacent image tiles in the stitching results of different methods as follows:

$$re_{i,j} = \sqrt{\left(\left|x_i - x_j\right| - x_s\right)^2 + \left(\left|y_i - y_j\right| - y_s\right)^2}$$

where $re_{i,j}$ is the relative offset error between the $i$-th and $j$-th image tiles (adjacent image tiles), ($x_i$, $y_i$) and ($x_j$, $y_j$) are the coordinates of the image tiles in the stitching result, and ($x_s$, $y_s$) is the coordinate of standard.

According to the “track lines” designed on the Stereo-seq chip, a template of stitched image can be obtained for each Stereo-seq dataset (the public dataset has no “track lines” that its template is not obtained). These templates are used to calculate the absolute offset error in the results of different methods as follows:

$$ae = \sum_{k=1}^{n} \sqrt{(xr_k - xt_k) + (yr_k - yt_k)}$$

where $ae$ is the absolute offset error, ($xr$, $yr$) and ($xt$, $yt$) are the coordinates of a marker point in the stitching result and template respectively, $k$ means the $k$-th marker point ($k$ = 1, 2, …, $n$), and there are total $n$ marker points obtained according to the “track lines”.

**Data analysis process and evaluation metrics in experiment of generating single-cell spatial gene expression profile on Stereo-seq mouse olfactory bulb dataset.**

The profiles are input into Stereopy (v6.0) to calculate the total gene counts and number of uniquely expressed genes through the quality control function. During the filtering process, the cells with fewer than 150 expressed genes and more than 5% mitochondrial genes are removed, and genes present in less than 3 cells are also removed. The profiles are then normalized using the "SCTransform" function. The differentially expressed genes are summarized by Principal Component Analysis (PCA) to reduce the data dimensionality. With these settings, we run the uniform manifold approximation and projection (UMAP) algorithm to obtain 2D data projections, followed by the "Leiden" clustering to identify all clusters within the dataset. The silhouette coefficient for evaluating clustering results is calculated as follows:

$$S(i) = \frac{b(i) - a(i)}{\max\{a(i), b(i)\}}$$

where, $S(i)$ is the silhouette coefficient, $a(i)$ indicates the average distance between the $i$-th sample and other samples in its cluster, and $b(i)$ is the average distance between the $i$-th sample and the samples in other clusters. Silhouette coefficient provides information on how similar a given cell is to other similar cells/bins (cohesion) in comparison with non-similar cells (separation). The moran's I for evaluating spatial correlation as follows:

$$I = \frac{n}{S_0} \frac{\sum_{i=1}^{n} \sum_{j=1}^{n} \omega_{i,j} z_i z_j)}{\sum_{i=1}^{n} z_i^2}$$

where, $I$ is the moran's I, $z_i$ is the deviation of the attribute of factor $i$ from its mean value, $w_{i,j}$ is the spatial weight between factors $i$ and $j$, $n$ is equal to the factor integration, and $S_0$ is the aggregation of all spatial weights.

**Data analysis process and evaluation metrics in experiment of dissecting the structural composition on Stereo-seq mouse brain dataset.**

The generated profiles are input into Stereopy (v6.0), the total gene counts and number of uniquely expressed genes are calculated, the cells with fewer than 200 expressed genes and more than 5% mitochondrial genes are removed, and the genes present in less than 3 cells are removed. The profiles are normalized using the "SCTransform" function. The differentially expressed genes are summarized using PCA and the 2D data projections are obtained by the UMAP algorithm, followed all clusters within the dataset are identified by the "Leiden" clustering.

**Additional experiments: CellBin can be applied to diverse tissue datasets and outperform other state-of-the-art methods.**

In addition to mouse brain and mouse olfactory bulb datasets, CellBin can be applied to more datasets from diverse tissues, such as mouse embryo, mouse tongue and mouse heart. We conduct an additional set of experiments on a public dataset to further compare the CellBin's stitching algorithm, MFWS, with competitive methods, ASHLAR and MIST. The dataset used is "Day2 Plate" (https://isg.nist.gov/deepzoomweb/data/referenceimagestitchingdata) provided by the MIST paper. The evaluation metrics used are provided by the supplementary documentation (https://static-content.springer.com/esm/art%3A10.1038%2Fs41598-017-04567-y/MediaObjects/41598_2017_4567_MOESM1_ESM.pdf) of the MIST paper, which are also used in the ASHLAR paper. According to the script corresponding to the evaluation metrics, the dataset is divided into three levels of test sets, include the complete dataset with all tiles (Level 3) and the two subsets with partial tiles intercepted from Level 3 dataset (Levels 1 and 2 respectively). We also compare CellBin with state-of-the-art methods, SCS and Baysor. SCS combines imaging data with sequencing data for nuclei segmentation to generate the spatial gene expression profiles, however, it lacks the ability of image stitching and registration. SCS requires the input of staining image that have been stitched and registered, with the corresponding spatial gene expression data. Although SCS can process Stereo-seq dataset with a large-field-of-view, it essentially splits a large staining image into multiple patches and processes them separately, thus, it requires a high computing resource and a long runtime. Baysor can generate spatial gene expression profiles without using staining images, that is, it takes only the spatial gene expression data as input. Baysor also requires a high computing resource and a long runtime. Using the same computational resources as running CellBin, both SCS and Baysor fail to produce results smoothly on the complete mouse brain and mouse olfactory bulb datasets. Issues such as out of memory, abnormal program interruptions, or no output within an acceptable time occur. To facilitate effective comparison, we extracted two subsets from the demo dataset (mouse brain) and manually labeled their nuclei masks as the ground truths. Subset 1 includes a 512pixels×512pixels nuclei staining image and its corresponding spatial gene expression data with 380,198 molecules. Subset 2 comprises a 512pixels×512pixels nuclei staining image and its corresponding spatial gene expression data with 416,135 molecules.

CellBin successfully and efficiently obtains single-cell spatial gene expression profiles on mouse embryo, mouse tongue and mouse heart datasets, and these profiles are used for clustering (Fig. S1a). Notably, the clustering results on the mouse embryo dataset exhibit clear separation of different organ outlines, which is highly consistent with the clustering results of a similar mouse embryo sample in the published Mouse Organogenesis Spatiotemporal Transcriptomic Atlas (MOSTA) (enter MOSTA web page https://db.cngb.org/stomics/mosta/, click “Spatial clustering” item and select “E16.5_E2S12.MOSTA.h5ad” as the File name). Some cell types in the mouse tongue and mouse heart datasets are also clearly distinguished. CellBin exhibits high efficiency and stability when applied to datasets of mouse embryos, which comprise multiple organs, as well as specific organ datasets such as the mouse brain, olfactory bulb, tongue, and heart. This provides strong evidence for the extensive applicability of CellBin across diverse tissue datasets.

From the results of stitching comparison experiment, there is no significant difference in the results of MFWS, ASHLAR, and MIST in terms of the area error metric (Fig. S1b, right column). In terms of the distance error metric, MFWS has a slight advantage over MIST and a significant advantage over ASHLAR on Level 1 dataset (Fig. S1b, top of left column). On Level 2 dataset, MFWS outperforms both ASHLAR and MIST (Fig. S1b, center of left column). On Level 3 dataset, MFWS performs better than MIST but worse than ASHLAR (Fig. S1b, bottom of left column). Overall, MFWS demonstrates competitive performance on the public dataset, comparable to MIST and ASHLAR.

On Subset 1, Baysor fails to output the results, while SCS and CellBin successfully output the results for analysis (Fig. S1c). The number of cells identified, number of clusters, silhouette coefficient, and Moran's I show minor differences between the two methods. However, SCS takes 33.76 times of runtime compared to CellBin. On Subset 2, Baysor, SCS, and CellBin all successfully obtain the results for analysis (Fig. S1d). Regarding the number of cells identified, SCS and CellBin show minimal differences, but Baysor identifies a significantly higher number of cells compared to the ground truth. There are also no obvious differences in the number of clusters, silhouette coefficient, and Moran's I among the three methods. In terms of runtime, SCS is 24 times of CellBin, and Baysor is 166.9 times of CellBin. The obtained silhouette coefficients and Moran's I values for all methods are unsatisfactory on both subsets, possibly due to limited information provided by the subsets, which result in ineffective identification of cell relationships and inaccurate clustering results. Since CellBin and SCS are image-based methods, the number of cells identified by them closely matched the ground truth within a reasonable range. With reference to the ground truth, it is evident that Baysor erroneously identifies a considerable number of cells. As a method relying on spatial gene expression information, Baysor places higher demands on the number of captured molecules. While Baysor has demonstrated promising results with MERFISH data, it faces challenges in maintaining this advantage when dealing with in situ capture-based data (such as Stereo-seq data). This is primarily because in whole-transcriptome data, issues like diffusion and non-aggregation of molecules are commonly observed. In terms of runtime, CellBin showed a clear advantage. Considering all these results, CellBin emerges as the most reliable and efficient method for handling large-field-of-view and high-resolution data compared to state-of-the-art methods.

**Information of different tissue datasets.**

Hardware configuration information for our experiments:

A server with 40-core CPU (Intel(R) Xeon(R) CPU E5-2650 v3 @ 2.30GHz), 128GB of RAM and 24 GB of GPU (NVIDIA GeForce RTX 3090), and at least 12 GB of disk storage for each dataset to save all results.

Dataset information:

Mouse brain with 131,990,020 molecules and 117 image tiles.

Mouse olfactory bulb with 37,288,344 molecules and 143 image tiles.

Mouse embryo with 261,470,876 molecules and 156 image tiles.

Mouse tongue with 29,079,667 molecules and 110 image tiles.

Mouse heart with 34,122,689 molecules and 120 image tiles.
