## Supplementary figures and images for "CellBin: a highly accurate single-cell gene expression processing pipeline for high-resolution spatial transcriptomics"

### Supplementary Fig. S1

#### Supplementary Fig. S1

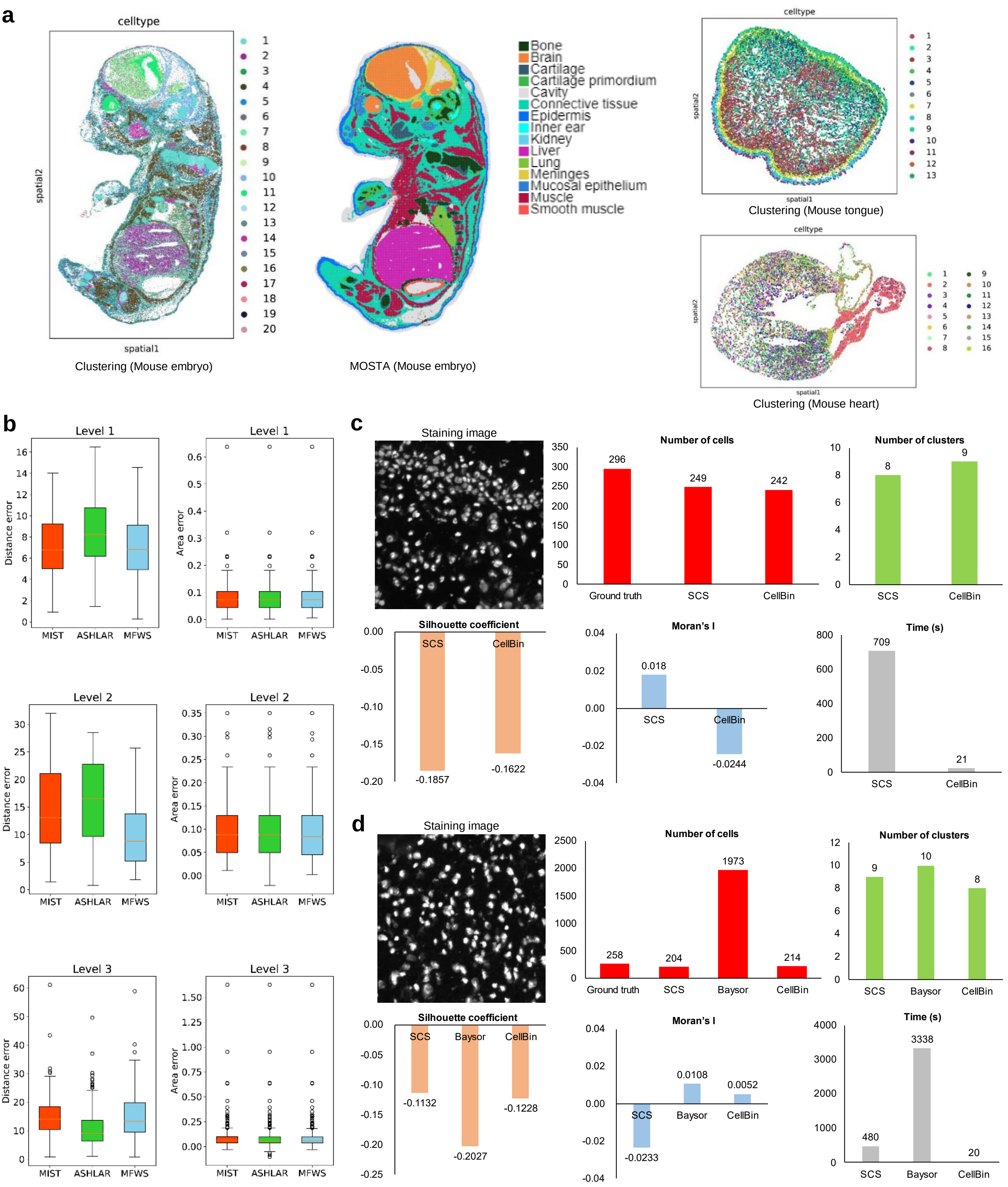
