## Supplementary Table S1 for "CellBin: a highly accurate single-cell gene expression processing pipeline for high-resolution spatial transcriptomics"

Table S1. Timing values of CellBin on different tissue datasets.

| Step | Mouse brain | Mouse olfactory bulb | Mouse embryo | Mouse tongue | Mouse heart |
|---|---|---|---|---|---|
| Image quality control | ~1m22s | ~1m12s | ~2m2s | ~1m39s | ~1m7s |
| Image stitching | ~1m38s | ~1m28s | ~2m16s | ~1m30s | ~1m23s |
| Image registration | ~3m32s | ~1m34s | ~7m29s | ~1m28s | ~2m54s |
| Tissue segmentation | ~10s | ~10s | ~15s | ~20s | ~10s |
| Nuclei segmentation | ~8m53s | ~8m26s | ~22m51s | ~6m54s | ~10m23s |
| Nuclei mask filtering | ~10s | ~14s | ~42s | ~14s | ~16s |
| Molecule labeling | ~1h5m27s | ~13m24s | ~8h35m13s | ~14m51s | ~27m20s |
